# ZMYND10 functions in a chaperone relay during axonemal dynein assembly

**DOI:** 10.1101/233718

**Authors:** Girish R Mali, Patricia Yeyati, Seiya Mizuno, Margaret A Keighren, Petra zur Lage, Amaya Garcia-Munoz, Atsuko Shimada, Hiroyuki Takeda, Frank Edlich, Satoru Takahashi, Alex von Kriegsheim, Andrew Jarman, Pleasantine Mill

**Affiliations:** MRC Human Genetics Unit, Institute of Genetics and Molecular Medicine, University of Edinburgh, Edinburgh, UK, EH4 2XU; Laboratory Animal Resource Centre, University of Tsukuba, Tsukuba, Japan, 305-8575; Centre for Integrative Physiology, University of Edinburgh, Edinburgh, UK, EH8 9XD; Systems Biology Ireland, University College Dublin, Dublin, Ireland; Department of Biological Sciences, University of Tokyo, Tokyo, Japan, 113-0033; Institute for Biochemistry and Molecular Biology, University of Freiburg, Freiburg, Germany, 79104; Department of Anatomy and Embryology, Faculty of Medicine, University of Tsukuba, Tsukuba, Japan, 305-8575; Edinburgh Cancer Research UK Centre, Institute of Genetics and Molecular Medicine, University of Edinburgh, Edinburgh, UK, EH4 2XU

## Abstract

Molecular chaperones promote the folding and macromolecular assembly of a diverse set of substrate ‘client’ proteins. How the ubiquitous chaperone machinery directs its activities towards a specific set of substrates and whether this selectivity could be targeted for therapeutic intervention is of intense research. Through the use of mouse genetics, imaging and quantitative proteomics we uncover that ZMYND10 is a novel co-chaperone for the FKBP8-HSP90 chaperone complex during the biosynthesis of axonemal dynein heavy chains required for cilia motility. In the absence of ZMYND10, defects in dynein heavy chains trigger broader dynein motor degradation. We show that FKBP8 inhibition phenocopies dynein motor instability in airway cells, and human disease-causing variants of ZMYND10 disrupt its ability to act as FKBP8-HSP90 co-chaperone. Our study indicates that the motile ciliopathy Primary Ciliary Dyskinesia (PCD) should be considered a cell-type specific protein-misfolding disease and opens the potential for rational drug design that could restore specificity to the ubiquitous chaperone apparatus towards dynein subunits.

## Introduction

Macromolecular motors of the dynein family power the essential beating of motile cilia/flagella. Motile cilia propel sperm cells, generate mucociliary clearance in airways, modulate nodal flow for embryonic left-right patterning and circulate cerebrospinal fluid inside the brain. Force-generating dynein motors are large molecular complexes visible by transmission electron microscopy (TEM), as ‘outer’ and ‘inner dynein arms’ (ODA, IDA) spaced at regular intervals along the microtubule axoneme. Each dynein motor consists of catalytic heavy chains (HC), intermediate chains (IC) and light chains (LC). In mammals, at least 4 ODA and 7 IDA subtypes exist, each containing different HCs^1^. Defective dyneins render cilia immotile, resulting in the severe congenital ciliopathy in humans termed Primary Ciliary Dyskinesia (PCD, OMIM: 242650). Understanding the molecular causes of PCD requires addressing the key question: how are complex molecular machines like the dyneins built during cilium biogenesis?

PCD causing mutations are most frequently detected in genes encoding structural ODA subunits, the intermediate chains (*DNAI1* and *DNAI2* ^2,3,4^), the redox light chain (*DNAL1*^5^) or the catalytic heavy chain (*DNAH5*^6^), all of which disrupt motor assembly and/or functions. Consequently, mutant multiciliated cells form cilia but these fail to move, lacking ODAs by TEM or immunofluorescence.

Several PCD causing mutations are also found in a newly discovered set of genes, the "dynein axonemal assembly factors" (DNAAFs), whose functions are poorly understood. DNAAFs are proposed to assist Heat Shock Protein (HSP) chaperones to promote subunit folding and cytoplasmic pre-assembly of dynein motors. DNAAFs are presumed to act as cilial-specific co-chaperones based on proteomic identification of interactions with both “client” dynein chains and canonical chaperones. Of the known assembly factors, KTU/DNAAF2 and DYX1C1/DNAAF4 have the most direct biochemical links to HSP90 and HSP70 chaperones, as well as ODA intermediate chain 2 (IC2) ^7,8^. High levels of homology between several DNAAFs (PIH1D3/DNAAF6, DNAAF2 and SPAG1) and key non-catalytic subunits (PIH1 and TAH1) of the well-known R2TP co-chaperone complex, further implicate these DNAAFs in chaperoning functions. Interactions between LRRC6, DNAAF1/LRRC50 and C21ORF59/Kurly were recently reported which, coupled with the phenotypic analysis of *Lrrc6* mutant mice, suggests that these assembly factors may primarily function in apical targeting/trafficking of dynein complexes ^9,10^. The functions of DNAAF5/HEATR2 which has no reported links to chaperones, remain elusive ^11^.

Dynein pre-assembly has been best characterised in *Chlamydomonas*. For ODAs, affinity purification confirmed all three HCs (HCs; α, β, and γ, each of ~500 kDa) and two ICs (IC1, 78 kDa; IC2, 69 kDa) are pre-assembled as a three headed complex and exist in a cytoplasmic pool prior to ciliary entry ^12-14^. This cytoplasmic pre-assembly pathway is highly conserved and exists in all ciliated eukaryotes ^15^. While it is clear that the aforementioned assembly factors aid axonemal dynein preassembly, their precise molecular functions within the pre-assembly pathway still remain largely unknown.

Previous studies had established a strong genetic link between loss of ZMYND10 and perturbations in dynein pre-assembly ^16^, however the putative molecular role of ZMYND10 as a DNAAF in this process remains unclear. In order to systematically probe the mammalian dynein pre-assembly pathway in greater molecular and cellular detail, we generated *Zmynd10* null mice by CRISPR gene editing. We used different motile ciliated lineages at distinct stages of differentiation from our mammalian mutant model to pinpoint the precise stage at which ZMYND10 functions during dynein pre-assembly.

Our protein interaction studies implicate a novel chaperone complex comprising of ZMYND10, FKBP8 and HSP90 in the maturation of dynein HC clients. We postulate that a chaperone-relay system comprising of several discrete chaperone complexes handles the folding and stability of distinct dynein subunits. Folding intermediates are handed off to successive complexes to promote stable interactions between subunits all the while preventing spurious interactions during cytoplasmic pre-assembly.

## Results

### Generation of a mammalian PCD model to characterize dynein assembly

We targeted exon 6 of mouse *Zmynd10* to target all predicted protein isoforms, with three guide RNA (gRNA) sequences for CRISPR genome editing and generated several founders with insertion, deletion and inversion mutations (**Figure 1A, Figure S1**). Null mutations from the different CRISPR guide RNAs gave identical phenotypes, confirming the phenotypes are due to loss of ZMYND10, as opposed to off-target effects. For detailed analysis, we focused on a −7bp deletion mutant line (*Zmynd10* c.695_701 p.Met178Ilefs*183), which results in a frame shift with premature termination. Generation of a null allele was verified by ZMYND10 immunoblotting of testes extracts (postnatal day 26, P26) and immunofluorescence of multiciliated ependymal cells and lung cryosections (**Figure 1B-D**).

**Figure 1.**
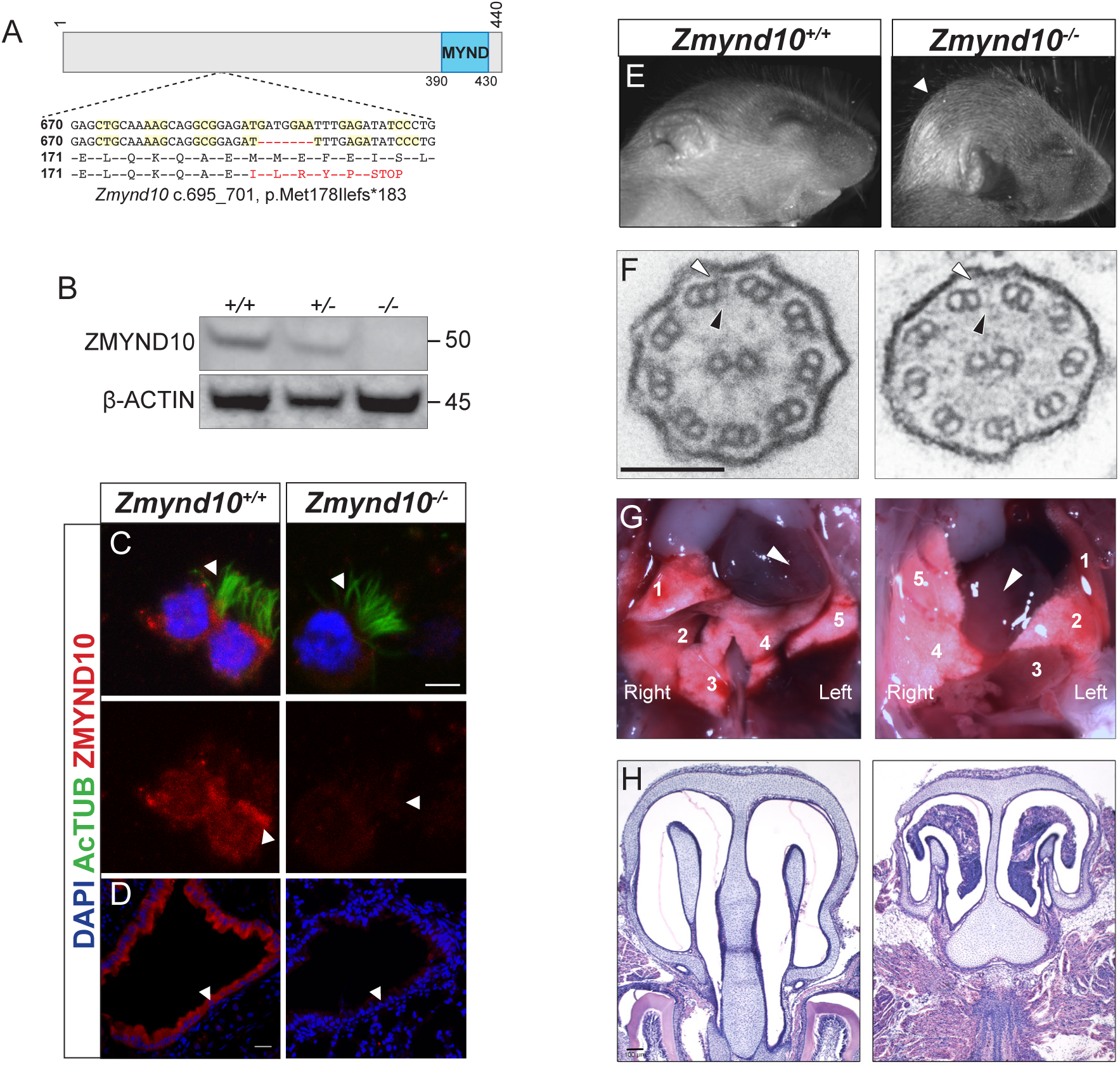
Loss of *Zmynd10* in mice results in a PCD phenotype. (A) Schematic illustrating the null allele generated by a −7bp CRISPR deletion in *Zmynd10* exon 6. (B) Immunoblots from testes extracts from postnatal day 26 (P26) control and mutant male mice show loss of ZMYND10. (C,D) Immunostaining for ZMYND10 reveals complete loss of signal in multiciliated ependymal cells (C) and lung cryo-sections (D). Multicilia are marked with acetylated α-tubulin (C). (E) Neonatal *Zmynd10* mutants display hydrocephaly; the white arrowhead points to doming of the head. See also Figure S2E. (F) TEM of trachea ciliary axonemes shows a lack of axonemal ODA (white arrowhead) and IDA (black arrowhead) dynein arms in mutants. (G) *Zmynd10* mutants display heterotaxy defects. Arrowheads denote direction of the heart’s position, which is reversed in the mutants; numbers denote lobes of the lungs (1,2 and 3 correspond to upper, mid and lower lobes of the right lung; 4 and 5 correspond to the upper and lower lobes of the left lung). See also Figure S2F. (H) H&E staining of coronal sections of nasal turbinates reveals mucopurulent plugs in mutants. Scale bars in (C)=5μm, in (D)=100μm in (E)=100nm.

*Zmynd10* mutant mice displayed several clinical features of PCD including heterotaxia, progressive hydrocephaly and chronic mucopurulent plugs in the upper airways, all features consistent with defects in ciliary motility (**Figure 1E-H**). This was directly confirmed by high-speed video microscopy of ependymal cells, where cilia of normal number and length were present but failed to move (**Movie 1,2,3,4, Figure S3**). Ultrastructure analysis of tracheal cilia axonemes revealed an absence of both outer and inner dynein arms (**Figure 1F**). The hydrocephaly phenotype was particularly pronounced on a C57BL6/J background and the majority of mutants died around weaning (P17-P21). On outbred backgrounds, male infertility and sperm immotility were also noted in homozygous mutant animals (Figure S2, Movie 5,6). These findings demonstrate that ZMYND10 functions are exclusively required in the motile ciliated cell lineages.

### Mis-assembled dynein motors are blocked from entering cilia and cleared in *Zmynd10* mutants

We analyzed expression of ODA components in different postnatal tissues by immunofluorescence and immunoblotting from *Zmynd10* mutants to assess whether ZMYND10 loss impacts ODA levels. In postnatal trachea and oviducts, total levels of the ODA HCs DNAH9 and DNAH5 were reduced by immunoblot (**Figure 2A,B**) and immunofluorescence (**Figure 2C,D**). In contrast to previous reports ^16^, no alteration in dynein transcripts were detected by RT-qPCRs on mutant oviduct total RNA, supporting the zinc-finger MYND domain of ZMYND10 plays cytoplasmic molecular scaffold functions other than a nuclear transcriptional role (**Figure 2E**). Critically, immunoblots of P7 *Zmynd10* mutant oviduct lysates, a stage corresponding to synchronized multicilial axonemal elongation ^17^, showed a laddering of DNAH5 products indicating post-translational destabilization of DNAH5 in the absence of ZMYND10 (**Figure 2F**).

**Figure 2.**
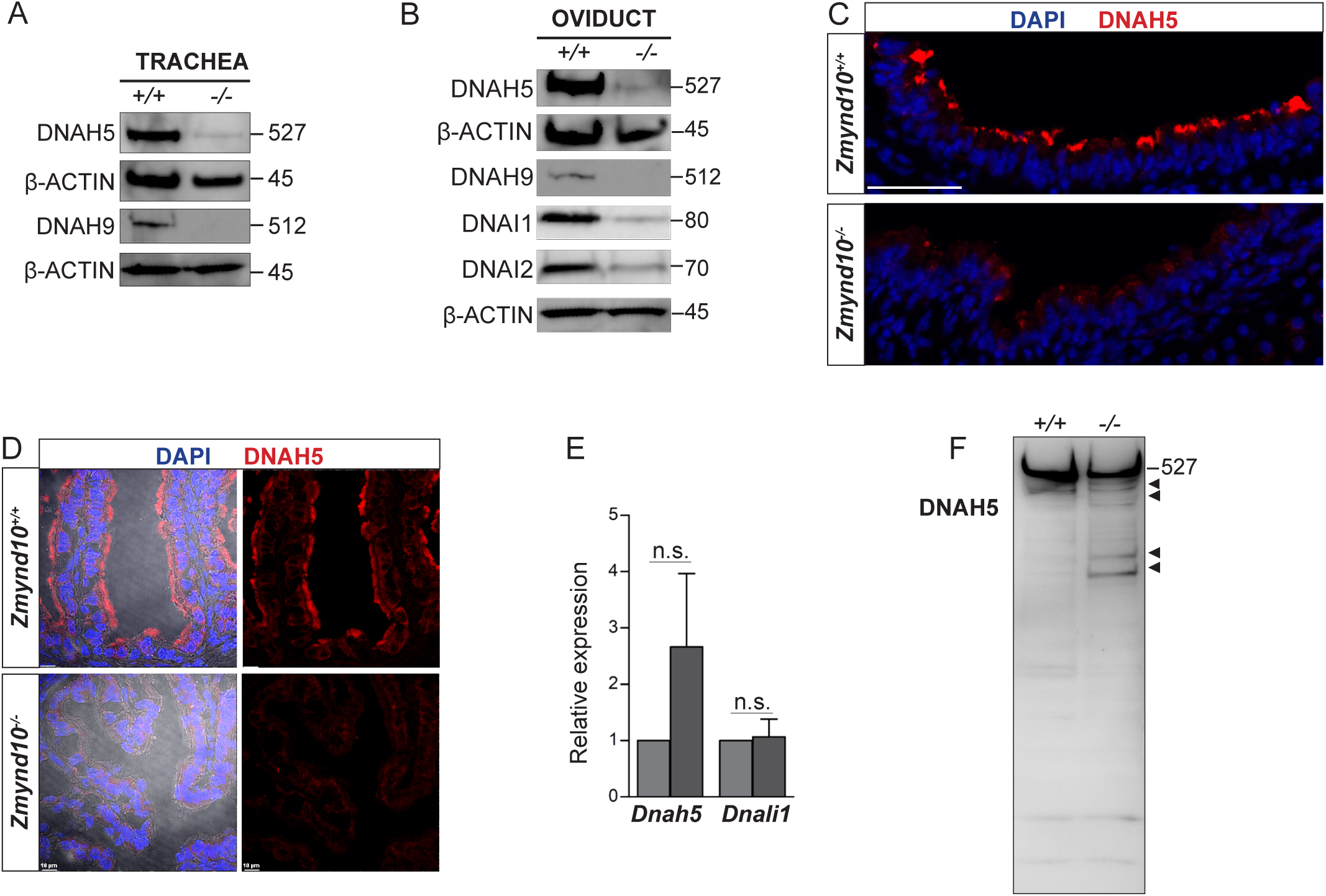
Global post-transcriptional destabilization of dyneins in *Zmynd10* mutant motile ciliated tissues. (A, B) Immunoblots of trachea P26 (A) and P30 oviduct (B) lysates show reduced abundance of both ODA HC-γ and HC-β (DNAH5 and DNAH9) in *Zmynd10* mutants compared to control littermates. (C, D) Immunofluorescence of trachea (C) or oviduct (D) tissue sections show loss of cilial DNAH5 staining as well as reduced total abundance in *Zmynd10* mutants. Scale bars in (C) & (D) = 100μm (E) No significant change by quantitative RT-PCR of levels of dynein transcripts (*Dnah5, Dnali1*) normalized to (*Tbp*) is detected in P12 *Zmynd10* mutant oviducts (n=3/genotype, dark grey *Zmynd10* mutants). (F) During early motile ciliogenesis, reduced levels and laddering consistent with degradative, misfolded intermediates of ODA HC-γ DNAH5 are detected in *Zmynd10* mutant oviducts (P7). These will be subsequently cleared as differentiation proceeds.

We previously reported distinct morphological and dynein staining characteristics observed among human multiciliated nasal brush biopsied cells depending on their maturity^11^. In cells isolated from nasal turbinates of control mice, ‘immature’ multiciliated cells were rounder with higher cytoplasmic dynein immunostaining and shorter cilia in keeping with cytoplasmic pre-assembly of the motility machinery. In contrast, ‘mature’ cells have long, organized arrays of cilia intensely stained for ODA subunits with a clear lack of cytoplasmic signal, suggesting that the motility machinery has translocated into cilia and stably integrated into the axonemal ultrastructure (**Figure 3A,B** upper panels). In *Zmynd10* mutants, no defects in ciliary length or number were observed (**Figure S3**) however outer or inner arm dyneins fail to incorporate into mature cilia axonemes. Importantly, no cytoplasmic accumulations were noted in ‘mature’ ciliated cells (**Figure 3A** lower panel). Surprisingly, strong dynein staining in the cytoplasm was observed in both ‘immature’ control and *Zmynd10* mutant cells, indicating ODA and IDA dynein subunit precursors are initially synthesized normally, further supporting that ZMYND10 loss does not impact their transcription or translation. Instead, loss of ZMYND10 leads to dyneins being robustly cleared when their pre-assembly is perturbed.

**Figure 3.**
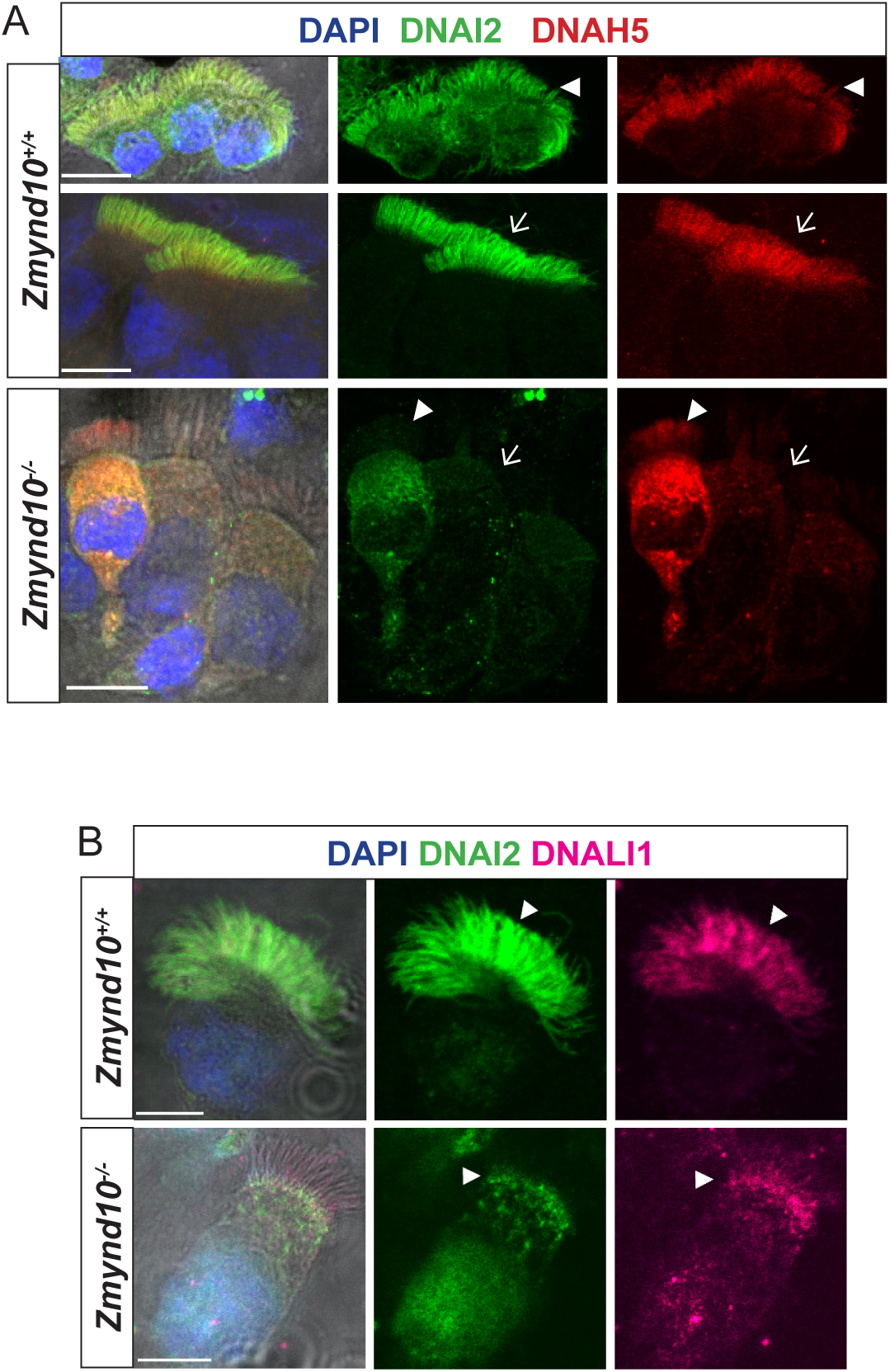
Loss of ZMYND10 perturbs sub-cellular distribution and levels of dynein complexes. Nasal brush immunofluorescence from *Zmynd10* mice shows both components of (A) outer arm and (B) inner arm dyneins are initially expressed in apical cytoplasm of immature mutant cells (arrowheads) but subsequently undergo ‘clearance’ in mature cells (lower panels, arrows), whilst all complexes exclusively translocate into cilia in control mature cells (upper panels, arrow). Scale bars =5μm

### ODA and IDA complexes are defective and unstable in the absence of ZMYND10

As cytoplasmic DNAI2 and DNAH5 were detected in *Zmynd10^-/-^* immature respiratory cells, suggesting that they were initially synthesized, we sought to verify if they were assembled into complexes using the *in situ* proximity ligation assay (PLA). In control immature human nasal brush epithelial cells, we detected PLA signals consistent with DNAI2 and DNAH5 existing in both cytoplasmic and axonemal complexes (**Figure 4A**). However, we detected a highly reduced number of PLA positive foci in nasal epithelial cells of P7 *Zmynd10*^-/-^ mice, with complexes restricted entirely to the cytoplasm in contrast to the strong axonemal staining observed in similarly staged controls (**Figure 4B**). To directly examine the interactions between ODA IC and HC subunits, we immunoprecipitated endogenous DNAI2 (IC2) from postnatal tracheal (P26) and oviduct (P7) extracts from *Zmynd10*^-/-^ animals. DNAI2 co-precipitated DNAI1 (IC1) at similar levels from both wild type and mutant P26 tracheal extracts (**Figure 4C**). This indicated that loss of ZMYND10 does not primarily impact IC subunit heterodimerization or stability during the assembly process. Importantly, we observed significantly reduced co-immunoprecipitation of DNAH5 by DNAI2 in P7 oviduct mutant extracts (mutant 0.55 vs wild type 1.1, normalized to total levels, Figure 4D,E). Moreover, we observed similar degradative bands (arrowheads) for DNAH5 in the mutant samples indicating that any DNAH5 that is incorporated may be poorly folded and unstable, in the absence of ZMYND10. We hypothesize that this reduced association between the two subunits is due to the HC subunit being in an assembly incompetent, unstable state such that any substandard complex would be targeted for subsequent degradation (**Figure 4F**).

**Figure 4.**
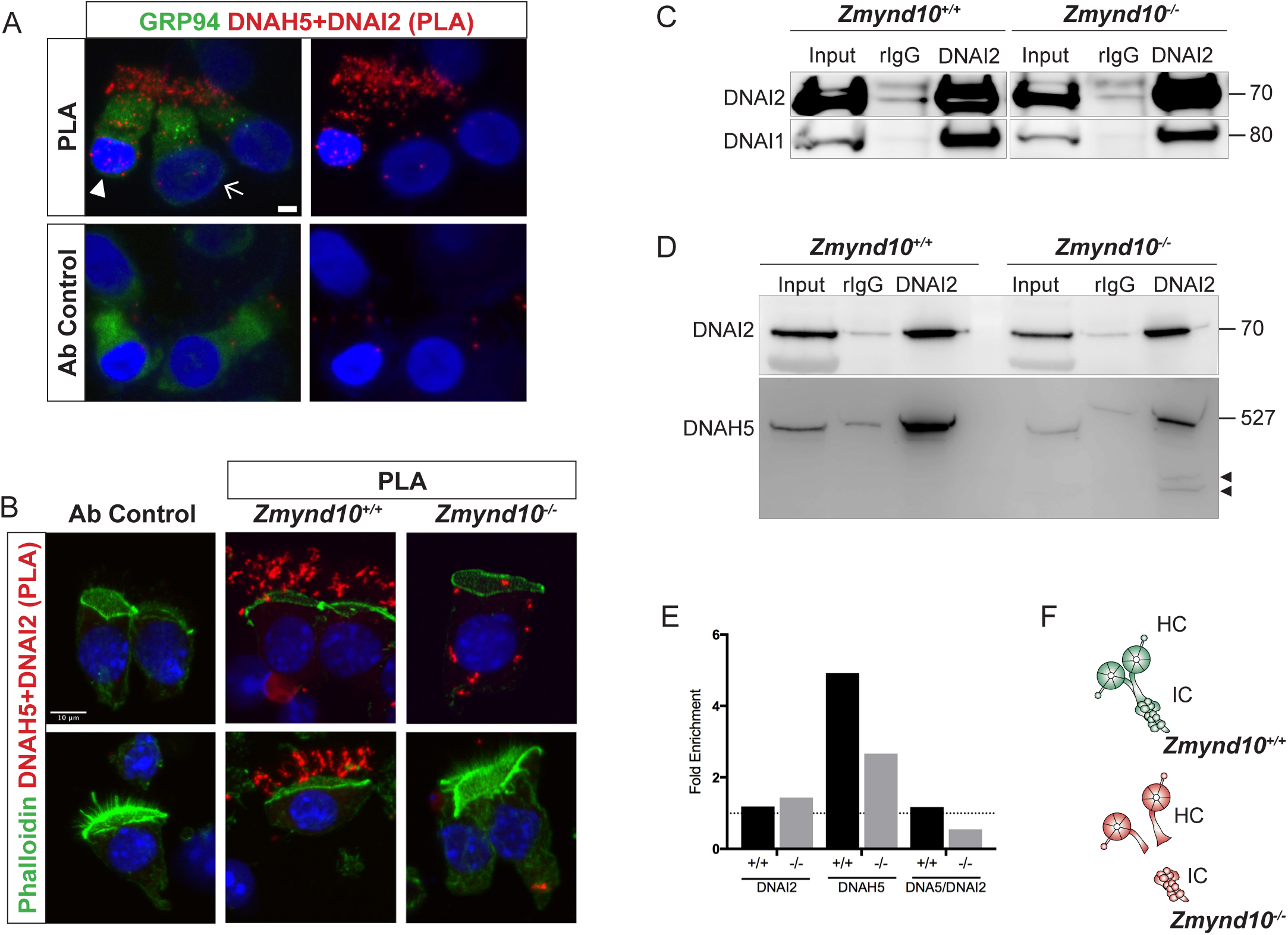
Sequential cytosolic assembly of outer arm dynein components occurs in mammalian motile ciliogenesis in a process requiring ZMYND10. (A,B) Proximity Ligation Assay (PLA) on human (A) or mouse P7 (B) nasal brush biopsies confirms that ODA subunits (mouse IC2/DNAI2 and HC-γ/DNAH5) are pre-assembled in the cytoplasm of mammalian multiciliated cells. Control (Ab control) sections were incubated with only DNAH5. Red spots denote individual ODA complexes (<40 nm) that appear as peri-nuclear foci in immature cells (arrowhead) and translocate to cilia in mature cells (arrow). In *Zmynd10* mutant cells, reduced number of foci are observed and restricted to the cytoplasm, highlighting defects in cytoplasmic assembly. GRP94 was used as a pan-cytosolic marker (A) or phalloidin for apical actin ring (B). Nuclei are stained with DAPI (blue). Scale bars in (A)=5μm and (B)=10μm (C) Immunoprecipitation of DNAI2 from P26 trachea extracts reveals no defects in DNAI2 association with its heterodimeric partner DNAI1 in *Zmynd10* mutants. (D) Immunoprecipitation of DNAI2 from P7 oviduct extracts show disruption in subsequent association between DNAI2 and DNAH5 in mutants compared to controls as quantified by intensities of the DNAH5 pull-down bands (E). Arrowheads show predicted degradative or misfolded intermediate of DNAH5 polypeptide in the mutants only. Numbers to the right of panels denote protein molecular weight in kDa. (E) Quantification of fold enrichment (from D: IP/input) for DNAI2 and DNAH5, as well as amount of DNAH5/DNAI2 complexes, normalized for differences in stability in input. (F) Schematic of axonemal ODA showing the intermediate chain heterodimers (IC) bind normally to heavy chains (HC) to form the entire motor complex in controls (green) and that this association is perturbed in mutants (red).

ZMYND10 loss also leads to absent IDA motors from human, fly and mouse cilia ^16^. To bypass the limitation of robust immunoreagents for IDA detection, we used label-free quantitative proteomics comparing postnatal testes extracts from P25 control and *Zmynd10* mutant littermates (**Figure 5A-C**). We hypothesized that synchronized spermiogenesis and flagellar extension at this stage would correspond with cytoplasmic pre-assembly of flagellar precursors. Whilst protein expression profiles were not different between mutant and controls for differentiation, meiosis and cell death markers (**Table S2**), the expression profile for the motility machinery showed specific and significant changes wherein almost all the dynein HCs (outer and inner) detected were reduced whilst the other axonemal dynein subunits were generally not significantly changed (ICs WDR78 and DNAI1). This is distinct from previous observations in *Chlamydomonas*, where loss of DNAAFs (DNAAF1, 2 and 3) impacting HC stability generally led to an aberrant cytosolic accumulation of IC subunits^18^, highlighting a key difference between the two model systems. Components of the radial spokes (RS) and dynein regulatory complex (DRC) were also unchanged (**Figure 5D**). Interestingly, several DNAAFs including the co-chaperones DNAAF4 and DNAAF6 were moderately but significantly up regulated in *Zmynd10* mutants suggestive of a proteostatic response to counter aberrant pre-assembly as it progresses.

**Figure 5.**
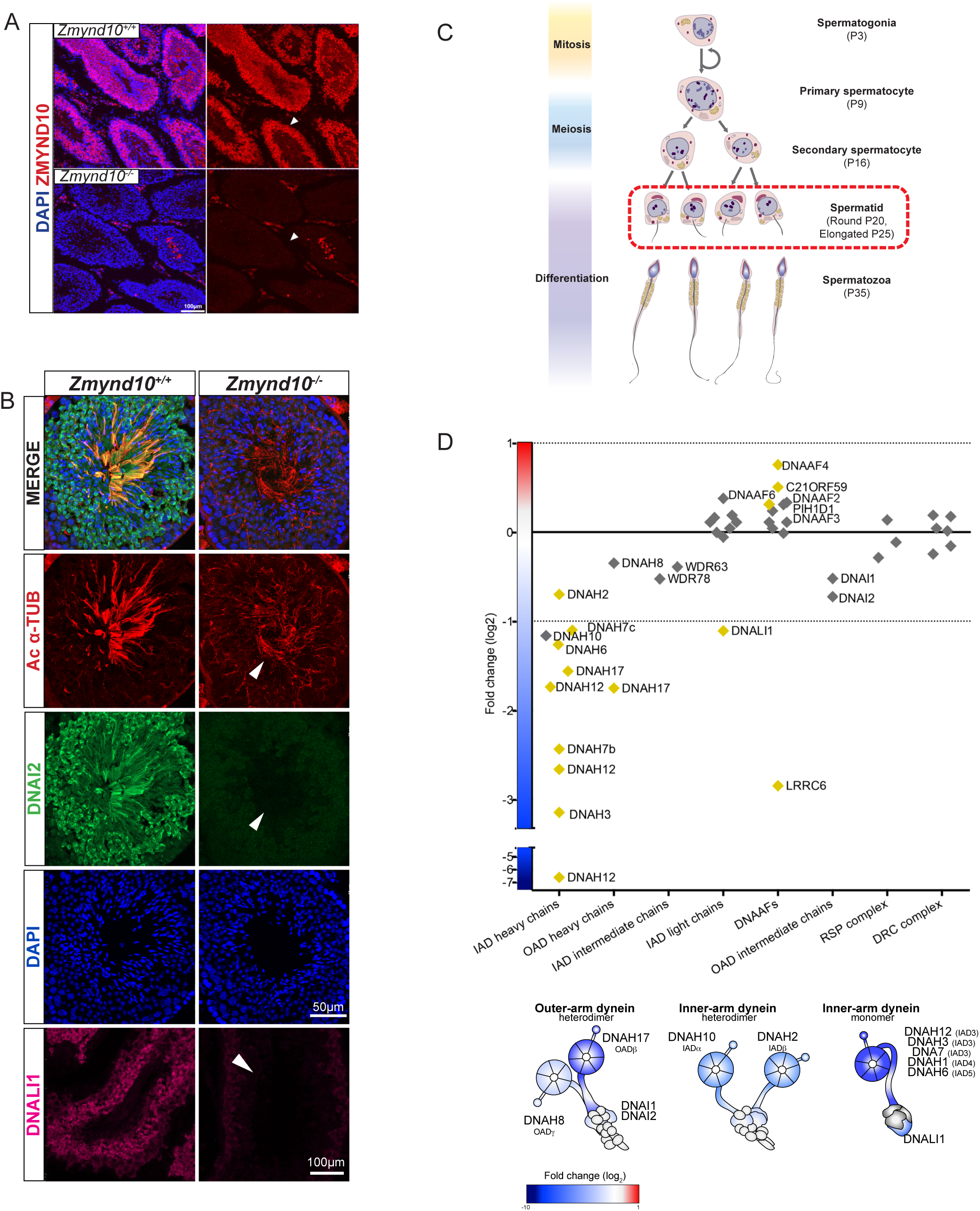
Early and specific defects in axonemal dynein heavy chain stability are observed in *Zmynd10* mutants during cytoplasmic assembly. (A, B) Immunofluorescence of ZMYND10 in control and mutant adult testes (P150) in asynchronous seminiferous tubules (arrowhead). (A) ZMYND10 is strongly expressed in primary spermatocytes and spermatids, where it is restricted to the cytoplasm and never in developing sperm tails. This staining is lost in mutants. Scale bar = 100μm. (B) Cross-section of similarly staged seminiferous tubules reveal similar developmental staging of sperm between control and mutants, but loss of DNAI2 (ODA) and DNALI1 (IDA) proteins from cytoplasm and axonemes (arrowhead) of mutant sperm. Scale bar = 50μm. (C) Schematic summarizing mouse spermatogenesis which is initially synchronized postnatally (stages shown to right), before occurring in asynchronous waves across seminiferous tubules. Axonemal dynein pre-assembly in the cytoplasm occurs in the spermatid stage. (D) Unbiased quantitative proteomics of control and mutant testes (elongated spermatid: P25) reveal that loss of ZMYND10 leads to a primary reduction in all dynein HC subunit abundance during cytoplasmic assembly, whilst other components are initially unaffected. Yellow squares highlight significantly changed hits between *Zmynd10* mutants and wild type littermates. (n=3/genotype). Schematic below highlights fold change of specific subunits on given dynein arms. Some heavy chains had multiple isoforms detected (*ie.* DNAH7). See also Supplemental Table 2.

### A ZMYND10-FKBP8-HSP90 complex mediates maturation of dynein heavy chains

To further understand how the loss of ZMYND10 results in instability of ODA HC subunits, we undertook a series of candidate protein interaction studies using two validated commercial ZMYND10 polyclonal antibodies. Firstly, we did not detect interactions with IFT-B proteins involved in ODA transport (**Figure S4**). Next, we investigated whether ZMYND10 associates with chaperones or other dynein assembly factors (**Figure 6A**), namely DNAAF2 which was shown to interact with DNAAF4, which also co-precipitates several chaperone proteins including the CCT3 subunit of the TriC chaperonin complex^8^ as well as the non-canonical co-chaperone DNAAF5, whose HEAT repeats have been proposed to function as scaffolds in multisubunit assembly^11,19^. We failed to detect interactions with HSP70, DNAAF2, CCT3 or DNAAF5 by endogenous ZMYND10 affinity purification from mouse testes or oviduct extracts (**Figure 6A,C**). Our findings suggest that ZMYND10 functions at a stage of dynein assembly distinct from these previously described cochaperone complexes. We were also unable to co-immunoprecipitate endogenous LRRC6, a protein that has been previously shown to associate with ZMYND10 by over-expression studies ^16^ Absence of strong association was confirmed by a reciprocal pull-down using endogenous LRRC6 as bait. We conclude that any interactions may be highly transient *in vivo* (**Figure 6B**).

**Figure 6.**
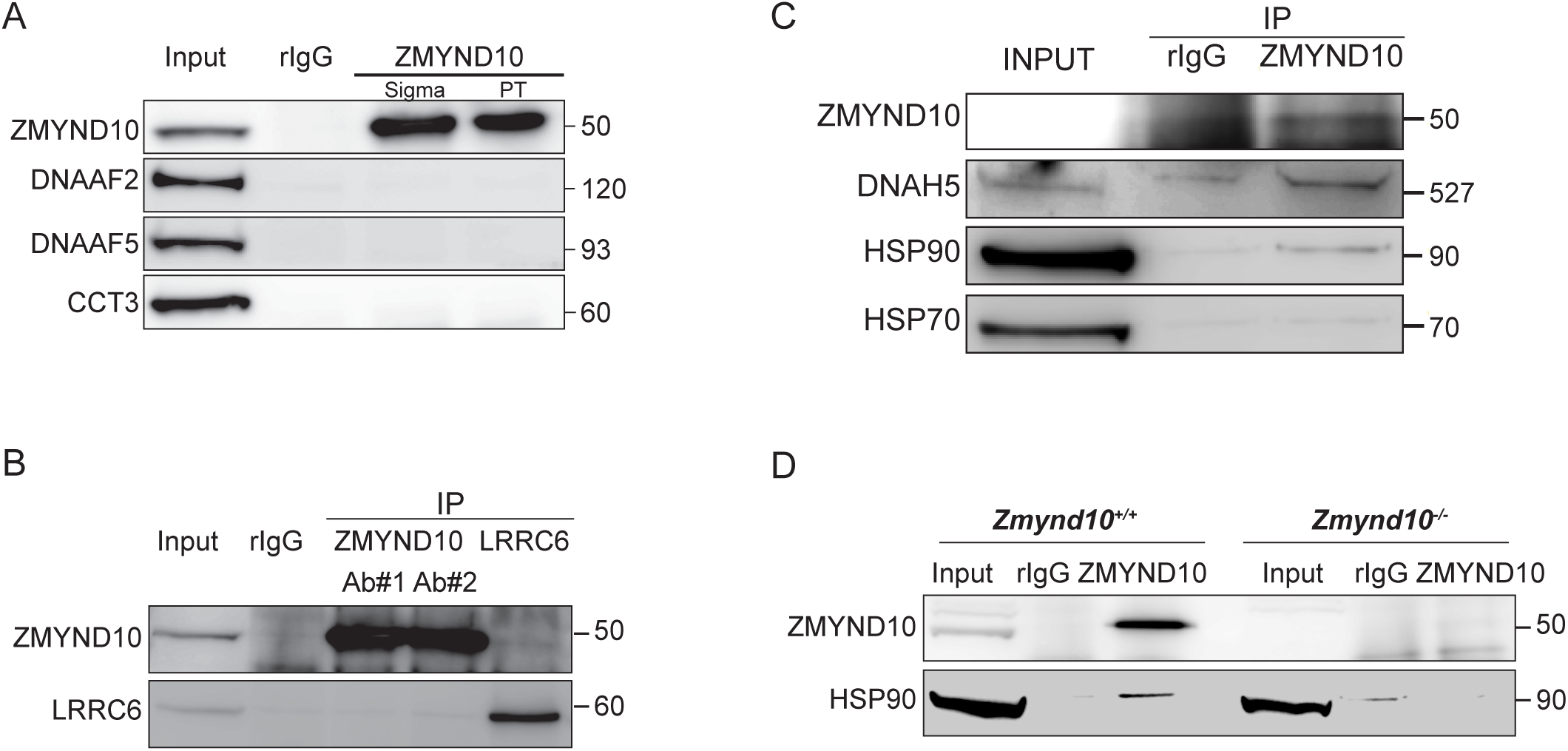
ZMYND10 interacts with a novel chaperone relay at a distinct stage of dynein heavy chain stability during cytoplasmic assembly. (A) Endogenous immunoprecipitation from P30 testes extracts using two ZMYND10 antibodies fail to immunoprecipitate DNAAF4 interactors DNAAF2 and CCT3, as well as DNAAF5. Protein molecular weights are displayed in kDa. (B) Failure to detect interaction *in situ* between LRRC6 and ZMYND10 in reciprocal endogenous immunoprecipitations in P30 control testes extracts, using rIgG as a control. (C) Endogenous ZMYND10 immunoprecipitation of client DNAH5 and chaperone HSP90, but not HSP70 from differentiating oviduct epithelial tissue (P7) using rIgG as a control. (D) Endogenous ZMYND10 immunoprecipitates HSP90 from P30 control testes extracts, but not *Zmynd10* mutant mice or rIgG controls.

We detected a specific interaction of ZMYND10 with HSP90, another major cytosolic chaperone implicated in dynein pre-assembly (**Figure 6C**). Our preliminary affinity purification mass-spectrometry (AP-MS) analyses to generate a ZMYND10 interactome (**Figure 7A, Table S1**) also revealed that ZMYND10 consistently co-precipitated the well characterised HSP90 co-chaperone

**Figure 7.**
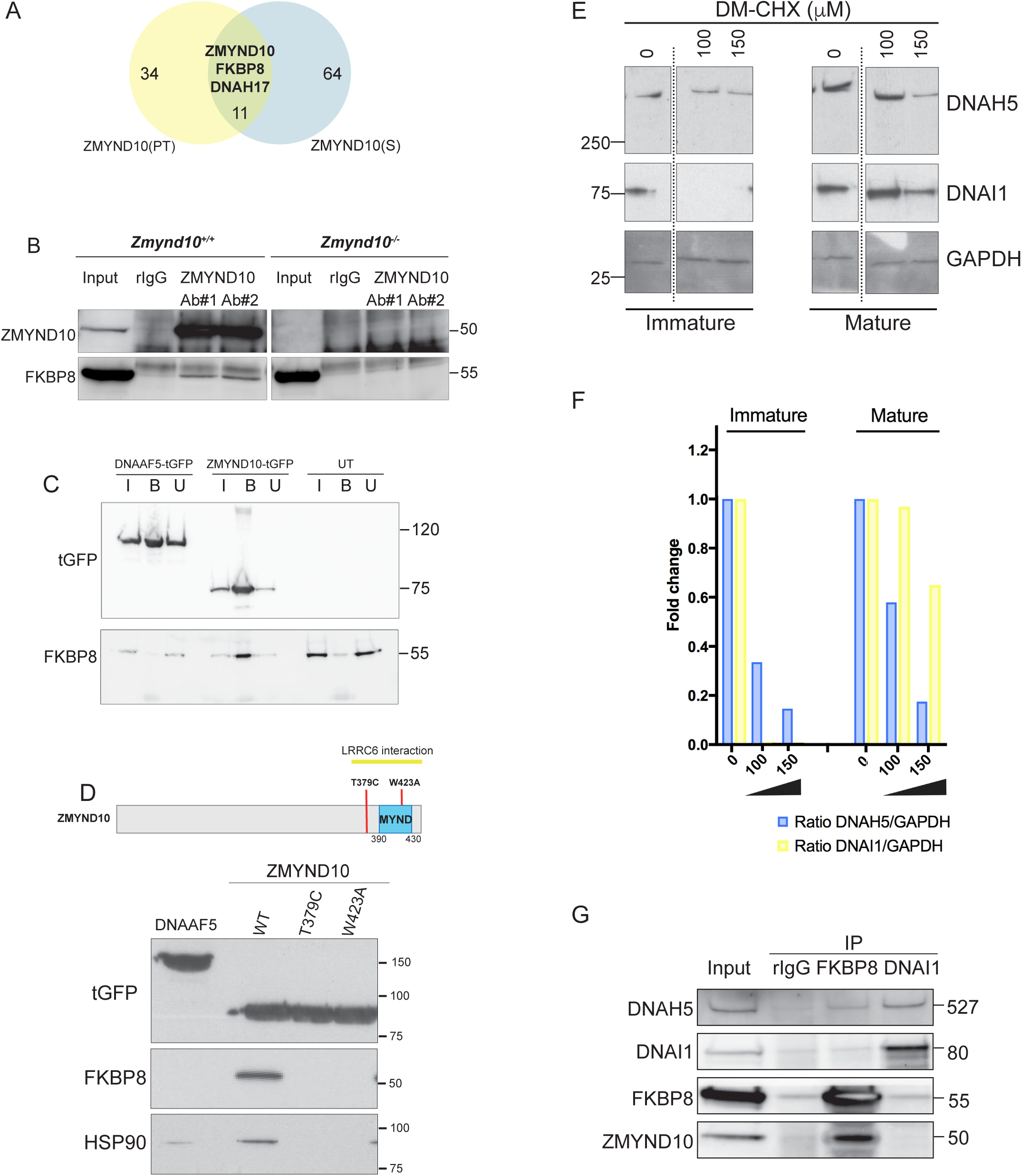
Ubiquitous FKBP8 actively participates in axonemal dynein heavy chain stability via its interaction with ZMYND10. (A) Summary schematic of mass spec analysis of interacting endogenous ZMYND10 immunoprecipitations from P25 mouse testes, overlapping between Sigma (S) and Proteintech (PT) polyclonal antibodies. See Table S1. (B) Endogenous ZMYND10 immunoprecipitation with two validated ZMYND10 antibodies from *Zmynd10* control and mutant P30 testes extracts show an interaction between ZMYND10 and FKBP8 in control samples. Mutant samples serve as pull-down controls. (C) Affinity purifications against turbo-GFP from HEK293T cells transiently transfected with DNAAF5-tGFP and ZMYND10-tGFP fusion proteins show that only ZMYND10, not DNAAF5, pulls-down endogenous FKBP8. Non-transfected cell lysates serve as pull-down controls. (D) Summary of mutations generated in ZMYND10-tGFP by site directed mutagenesis, to disrupt the binding interface for FKBP8 (middle panel). Input of transiently transfected RPE-1 cells with constructs, verifying expression (top panel). Affinity purification against turboGFP shows mutations interrupt FKBP8 and HSP90 binding. (E) Human tracheal respiratory epithelial cultures (MucilAir) before ciliation (D20) or full ciliated (D60), were cultured for 24 hours in control (vehicle only) or DM-CHX (concentrations indicated in μM) before harvesting protein extracts. Immature cultures (D20) were more sensitive to effects of specific PPIase inhibitor DM-CHX, destabilizing dynein components, whilst mature cultures were minimally affected. (F) Quantification of band intensities DNAH5 (blue) or DNAI1 (yellow) from (E) were normalized to loading control, and plotted as a fold change after 24 hours. (G) Endogenous FKBP8 and DNAI1 immunoprecipitations in differentiating human tracheal epithelial cultures (D17) both show binding of client DNAH5, the rest of the complexes are distinct, suggesting they act at sequential steps of assembly. Protein molecular weights displayed in KDa.

FKBP8 (Immunophilin FK506-binding protein (FKBP) family member; Uniprot: <u>O35465</u>), ^20^ as well as the ODA HC DNAH17. We confirmed the FKBP8 interaction with an endogenous IP strategy in control versus mutant samples (**Figure 7B**) as well as an additional tagged immunoprecipitation strategy for ZMYND10 (**Figure 7C**). Direct interaction between the N-terminus of FKBP8 and MYND domains of ZMYND6/PHD2 ^21^ and ZMYND20/ANKMY2 ^22^ have been reported. To map the interaction interface between FKBP8 and ZMYND10, we generated two point mutations in the MYND domain predicted to disrupt ZMYND10 function (**Figure 7D**). The W423A mutation is predicted to functionally disrupt one of two Zn^2+^-fingers in the MYND domain. Located just before the MYND domain, the T379C PCD patient mutation ^16^ failed to disrupt binding to LRRC6, suggesting some other underlying pathogenic mechanism exists for this mutation, one which we hypothesize could involve FKBP8. Affinity purification of ZMYND10-turboGFP variants from HEK293T cells revealed the point mutations abolish endogenous FKBP8 binding. These results indicate that the interaction interface for FKBP8 extends beyond the MYND domain of ZMYND10, consistent with recent deletion studies described in *medaka* capable of functional rescue ^23^.

FKBP8 is a peptidyl-prolyl isomerase (PPIase), which catalyzes cis-trans isomerization of proline peptide groups and is one of the rate-determining steps in protein folding. To test whether its PPIase activity is critical for stabilization of dynein HCs, we treated immature day 17 and mature day 60 (D17 and D60 in culture) human tracheal epithelial cells with a specific PPIase inhibitor DM-CHX ^24^ for 24 hours and assayed extracts for stability of ODA subunits by immunoblot. Immature cultures were very sensitive to FKBP8 inhibition, where levels of cytoplasmic DNAH5 were reduced to ~10% of control levels after DM-CHX (150μM). In mature cells, fully assembled complexes within cilial axonemes were less sensitive to DM-CHX treatment (**Figure 7E,F**). Surprisingly, a very striking destabilization of DNAI1 was also observed in immature cultures under these conditions. This supports the possibility that a transient requirement of the PPIase activity of FKBP8 is necessary for folding and/or stability of axonemal dyneins in the cytoplasm.

To directly test for ZMYND10 dependent chaperone-client associations, we immunoprecipitated endogenous FKBP8 and ZMYND10 from differentiating human tracheal epithelial cells (D17) and oviducts (P7); both these stages correspond to active cytoplasmic dynein pre-assembly. We detected associations of the client protein DNAH5 with both ZMYND10 and FKBP8 (**Figure 7G**). In the case of ZMYND10, it associated with HSP90 but not with HSP70 at a similar stage of cellular differentiation (Figure 6C). These findings show an association between a mammalian dynein assembly factor and a client dynein heavy chain *in vivo*. Additionally, taking step-wise ODA macromolecular assembly into account, we found that whilst DNAI1 co-immunoprecipitated DNAH5, it did not immunoprecipitate either ZMYND10 or FKBP8 (**Figure 7G**). This suggests that the DNAI1-DNAH5 interaction occurs in a complex that is distinct and downstream from the FKBP8-DNAH5-ZMYND10 complex, as supported by our DM-CHX experiments in which both subunits are destabilized after 24 hours drug treatment.

In summary, we propose that a complex or series of complexes of the cilial-specific ZMYND10 adaptor, the ubiquitous FKBP8 co-chaperone and chaperone HSP90 direct their activities towards client axonemal dynein heavy chains including DNAH5 to promote their maturation and stability. This ‘folding/maturation’ step is essential for the HCs to subsequently form strong associations with other dynein subunits as observed between DNAH5 and the IC1-IC2 heterodimer and for productive assembly to proceed.

## Discussion

Motile cilia are highly complex structures comprising of hundreds of mega-Dalton scale molecular complexes. Axonemal dynein motors represent the largest and most complex of such motile ciliary components. Their coordinated transcription, synthesis, assembly and transport is critically linked to ciliary function and the cell appears to have evolved a dedicated chaperone relay system involving multiple assembly and transport factors to execute distinct steps for their pre-assembly.

We show that ZMYND10 co-operates with the ubiquitous co-chaperone FKBP8 and chaperone HSP90 to mediate a key step in the pre-assembly pathway i.e. the maturation of the dynein heavy chain subunits. Using multiple ciliated tissues from CRISPR mouse models, we observed reduced protein abundances for ODA HCs, DNAH5 and DNAH9 in mature multiciliated cells, without effect on transcript levels. The observation of partially/mis-folded degradative intermediates of DNAH5 in *Zmynd10* mutant extracts further strengthens our hypothesis that loss of ZMYND10 severely affects HC post-transcriptional stability. Endogenous pull-down assays reveal that this unstable intermediate of DNAH5 is unable to fully associate with the IC1/2 heterodimer, which does form in *Zmynd10* mutants. Finally, inhibition of the PPIase activity of FKBP8 destabilizes wild type cytoplasmic dynein assemblies phenocopying what is observed in *Zmynd10* mutants, whilst PCD mutations of *ZMYND10* impair its ability to interact with the FKBP8-HSP90 chaperone system providing a molecular explanation for a previously unresolved disease-causing variant^16^. We propose that the aberrant HC-IC association and/or the misfolded HC polypeptides trigger a robust proteostatic response leading to clearance of non-functional ODA complexes to mitigate cellular protein stress.

Our unbiased label-free quantitative proteomics shows that ZMYND10 loss also specifically impacts IDA HC stability whilst other subunits, assembly factors or structures remain unaffected. This is distinct from the response seen in PCD models specifically affecting ODAs wherein complete lack of or misfolding of a single heavy chain in the case of DNAH5 results in a very specific and limited loss of outer dynein arms only: in *DNAH5* patients, DNALI1 is still found in the ciliary axonemes and IDAs visible by TEM^4^,

Our systematic protein-interaction studies are the first to begin mapping mammalian DNAAF functions under physiological conditions. Protein interaction studies and/or homology modeling for DNAAF2, DNAAF4, DNAAF6 and SPAG1 ^7,8,25,26^ place these assembly factors within a cilial-specific configuration of the HSP90 co-chaperone R2TP complex. DNAAF1, LRRC6 and C21ORF59 putatively function in another separate reported complex (Jaffe et al, 2016). Immunoprecipitation studies found no associations between ZMYND10 and several of these DNAAFs ^16^ highlighting a distinct function.

Our protein interaction and mutational studies define a novel ZMYND10-FKBP8-HSP90 complex functioning in dynein pre-assembly. In a motile cilia context, we describe a new function for the well-studied FKBP8-HSP90 chaperone complex ^20,21,27-32^. The role of this complex in ER-associated protein folding and maturation is well documented such as in folding of the CFTR protein on the cytosolic face of the ER ^30,31^. Our small molecule DM-CHX studies further highlight the requirement of FKBP8 PPIase activity for stabilizing dynein subunits at stages when cytoplasmic assembly is operating at its peak. Unlike *Zmynd10* mouse mutants, which display specific phenotypes consistent with motile cilia defects, *Fkbp8* mutant mice die shortly after birth from neural tube closure defects, likely due to both FKBP8 and HSP90 being ubiquitously expressed, whereas ZMYND10 is specifically expressed in multiciliated cell types. Taken together, we propose that in motile ciliated cells, ZMYND10 specifically directs the chaperone activities of the ubiquitous FKBP8-HSP90 complex towards stabilizing dynein HCs.

The strong interactions with ER resident FKBP8 and our PLA data together address a long-standing question of where dynein pre-assembly occurs within the cytoplasm. We observe that there is a transient wave of dynein subunits, which progressively traffic from the cytoplasm into nascent axonemes. When this process fails in mutants, like *Zmynd10*, subunits are cleared from the cytoplasm. We suggest that at this early stage of pre-assembly, dynein heavy chain synthesis could be localized to intracellular membranes such as the cytosolic face of the ER, which is a primary site for protein translation with access to a host of chaperones, including ER-resident FKBP8. Future work should be directed to further pinpoint where different assembly steps occur within the cytoplasm, such as the recently described dynamic cytoplasmic puncta {^33,34^.

Despite the molecular diversity in DNAAFs, a unifying theme is their participation in an emerging dynein assembly chaperone network specific to motile ciliated cells. Recent reports have highlighted the importance of PIH-domain containing DNAAFs (DNAAF2 and DNAAF6), TPR-domain containing SPAG1 and DNAAF4 as well as Reptin (RUVBL2) and Pontin (RUVBL1) as regulators ^7,8,33,35^ of cilia motility. All of these factors bear homology to or form part of the multi-functional R2TP complex (RUVBL1, RUVBL2, TAH1, PIH1). The role of R2TP as a co-chaperone of HSP90 in the multimeric assembly of snoRNPs and RNA Polymerase II is well established ^36,37^. Direct biochemical links have been reported between DNAAF2, DNAAF4, DNAAF6 and the ODA-IC2 subunit ^7,8,35^ Additionally, the IC1-IC2 heterodimer is specifically destabilized in DNAAF6/PIH1D3, RUVBL1 and RUVBL2 mutants ^35,33^. Critically, IC1-IC2 complex formation is not affected in *Zmynd10* mutants. Instead, our interaction studies show that the association between FKBP8, DNAH5 and ZMYND10 occurs separately to the association between DNAH5 and DNAI1/DNAI2. We suggest that an R2TP-like complex may function to predominantly stabilize the IC1-IC2 heterodimer in motile ciliated cells to serve as a platform for further subunit assembly. Based on our findings we would place ZMYND10 at a distinct node in the dynein assembly network, one that is critical for the post-translational maturation of dynein heavy chain subunits.

Taken together, we propose a revised model of the dynein preassembly pathway (Figure 8). Firstly, ZMYND10 acts a novel co-chaperone of the ubiquitous FKBP8-HSP90 chaperone complex for dynein HC subunit maturation. Mature, assembly competent HCs are then handed-off to a subsequent chaperone complex, likely the R2TP complex to allow for stable association with the IC1/2 complex, in a ZMYND10-independent step. Working together, this chaperone relay ensures efficient assembly of functional dynein complexes for subsequent ciliary targeting. Our work on ZMYND10 represents a paradigm shift in our understanding of PCD pathogenesis. We propose that the motile ciliopathy Primary Ciliary Dyskinesia (PCD) should be considered a cell-type specific protein misfolding disease, which may be amenable to therapy by modulation of the cellular proteostasis network.

**Figure 8.**
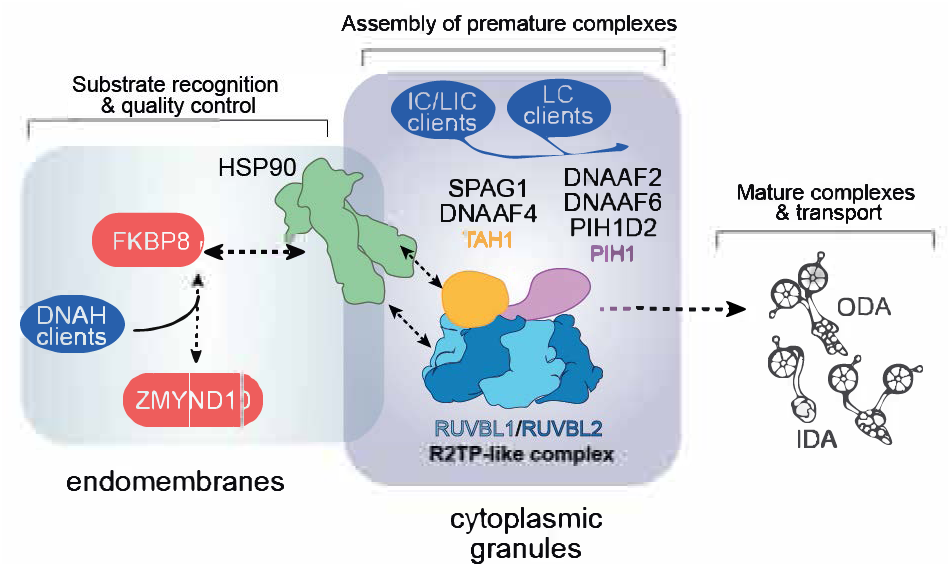
ZMYND10 specifies dynein heavy chains as clients in a chaperone relay during dynein pre-assembly. During cytoplasmic dynein arm assembly, ZMYND10 interaction with co-chaperone PPIase FKBP8 and HSP90 is required for the stabilization and folding of dynein heavy chains, as shown for DNAH5. We propose a specialized R2TP-like complex functions in the parallel assembly of heterodimeric DNAI1/2 complexes via the PIH-domains (DNAAF6, DNAAF2, PIH1D2), which may bind to the AAA+ATPases RUVBL1 and RUVBL2, and the TPR-domain containing SPAG1 and DNAAF4 which could in turn bind HSP90 via its MEEVD domain. This operates in parallel as DNAI1/2 heterodimers are detected in *Zmynd10* mutants, however these are likely degraded if fully functional mature complexes (with heavy chains) cannot be assembled. This chaperone-relay system comprises several discrete chaperone complexes overseeing the folding and stability of discrete dynein subunits. Folding intermediates are handed off to successive complexes to promote stable interactions between subunits all the while preventing spurious interactions. Stable dynein complexes, once formed are targeted to cilia via transport adaptors and intraflagellar transport (IFT).

## Materials and methods

### Generation of CRISPR mouse mutants

CAS9-mediated gene editing was used to generate mutant mice for *Zmynd10* (ENSEMBL:ENSMUSG00000010044) using three (guide) gRNAs each targeting “critical” exon 6. Guide RNA sequences were cloned into a pX330 vector (Addgene:#42230)^38^ and efficacy was first validated using a split GFP assay in HEK293 cells (Addgene: #50716)^39^. Pronuclear injections of 5ng/μl of purified plasmid DNA of pX330 constructs were injected into fertilized C57BL/6J eggs, which were cultured overnight until the two-cell stage before transferring to pseudopregnant females. PCR based screening, Sanger sequencing and characterization of genetic mutations of founder animals (F0) was performed. A genotyping was developed using a restriction digest of a PCR product for the −7bp deletion line used in this study. Animals were maintained in SPF environment and studies carried out under the guidance issued by the Medical Research Council in “Responsibility in the Use of Animals in Medical Research” (July 1993) and licensed by the Home Office under the Animals (Scientific Procedures) Act 1986.

### Cytology, Histology and TEM

Motile multiciliated ependymal cells were obtained from mouse brains (>P7) using a published protocol^40^. Mouse respiratory epithelial cells were obtained by exposing the nasal septum and scraping cells off the epithelium with an interdental brush (TePe, 0.8mm ExtraSoft) followed by resuspension in DMEM (isolated from animals at P9, P14 or P29). Cells were spread on superfrost slides, air-dried and processed for immunofluorescence. This was modified for proximity ligation assay (PLA) where cells were resuspended in PBS, then fixed 4%PFA/3.7% sucrose/PBS for 30 minutes on ice and cytospun onto Superfrost slides. Human respiratory epithelial cells by brush biopsying the nasal epithelium of healthy human donors or P7 neonatal mice were processed for proximity ligation assay using a Duolink PLA starter kit (DUO92101, Sigma-Aldrich), as per the manufacturer’s instructions following PFA fixation and 0.25%Triton-X100/TBS permeabilization 10 minutes. Alexa-488 phalloidin (Thermo Fischer) or rat anti-GRP94 (Thermo Fischer) counterstaining was done post-PLA protocol, prior to mounting in Duolink^®^ In Situ Mounting Medium with DAPI (Sigma Aldrich). Trachea, testes and oviducts were dissected and immersion fixed in 4% paraformaldehyde (from 16% solution, Thermo Fischer) overnight and cryosectioned for immunofluorescence^11^. Nasal turbinates were similarly fixed and processed for paraffin sectioning stained with H&E to reveal mucus plugs. Epidydymal spermatozoa were isolated by dissecting the cauda and caput regions of the epididymides in M2 media (Life Technologies), spread onto superfrost slides and air-dried followed by fixation and permeabilisation for immunofluorescence, as previously described^11^. For counting, sperm from the cauda epididymides were immobilized by diluting in H2O and counts were performed using a haemocytometer. For transmission electron microscopy, trachea tissue samples were dissected into PBS and immersion fixed in 2% PFA/2.5% glutaraldehyde (Sigma-Aldrich)/0.2M Sodium Cacodylate Buffer pH7.4 with 0.04% CaCl_2_^41^. Samples were cut into semi-thin and ultrathin sections and imaged by transmission electron microscopy (EM Services, Newcastle University Medical School).

### Live brain sectioning and high-speed videomicroscopy of ependymal cilia

Whole brains were isolated from neonatal mice in ice cold PBS and kept on ice. Brains were mounted vertically along the caudo-rostral axis on a petri dish and embedded in low melting point agarose (Thermo Scientific). 400μm thick vibratome sections of live brain tissue were obtained and floated onto wells of a glass bottom multiwell plate (Greiner Sensoplates cat.662892) containing DMEM and maintained at 37°C and 5% CO_2_. Sections were imaged on a Nikon macroscope to visualize dilated lateral ventricles. Motile cilia beating along the surfaces of the lateral walls were visualized and motility was recorded using a high-speed videomicroscopy Andor CCD camera attached to a confocal capture set-up.

### Immunoprecipitations (IP) and immunoblots

Endogenous immunoprecipitations were performed using protein extracts either from multiciliated cell cultures or motile ciliated tissues lysed under mild lysis conditions (50 mM Tris-HCl (pH 7.5), 100 mM NaCl, 10% Glycerol, 0.5 mM EDTA, 0.5% IGEPAL, 0.15% Triton-X 100 and Halt Protease Inhibitor Single use cocktail EDTA free (Thermo Fischer)). Modifications for detecting HSP90 interactions, included using sodium molybdate (Sigma-Aldrich) was included in the IP buffer. To detect interactions between ODA subunits, a DNAI2 antibody (Abnova) was used as a bait to enrich DNAI2 containing complexes from mouse trachea and oviduct lysates. Immunoblotting was performed using DNAI1 (Proteintech) and DNAH5 (M. Takeda). For ZMYND10 interaction studies, extracts from whole testes (P26) and differentiating ependymal primary cultures were used. Endogenous ZMYND10 containing complexes were pulled out using two validated ZMYND10 polyclonal antibodies (Sigma, Proteintech). Immunoblotting was performed using an HSP90 antibody (Santa Cruz). For human samples, endogenous FKBP8, DNAI1 and DNAI2 pulldowns were perfomed on lysates from normal human airway epithelial cells (MucilAir, Epithelix Sarl) grown at air-liquid interface for 17-19 days (immature cells). Antibodies for FKBP8 (Proteintech), DNAI1 (Proteintech) and DNAI2 (Abnova) were used as baits and antibodies for DNAH5 (Sigma) and ZMYND10 (Proteintech) were used to detect these interactors. An isotype-matched IgG rabbit polyclonal antibody (GFP: sc-8334, Santa Cruz) was used as control. In all pulldown experiments, immunocomplexes were concentrated onto Protein G magnetic beads (PureProteome, Millipore). Following washes, immunocomplexes were eluted off the beads and resolved by SDS-PAGE for immunoblotting. Alternatively, beads were processed for on-bead tryptic digestion and mass-spectrometric analysis. For overexpression pulldowns, transient transfection (Lipofectamine2000) of mouse *Dnaaf5-tGFP* (Origene-MG221395) and *Zmynd10-tGFP* (Origene, MG207003) into either HEK 293T or RPE1 cells. Site-directed mutagenesis was performed using two complementary PCR primers containing the desired nucleotide changes (PrimerX tool) to amplify *Zmynd10-tGFP* with proof reading DNA polymerase (Agilent II), followed by DpnI digestion, E coli transformation and sequencing of the thus recovered plasmids. Primer sequences available upon request. Subsequent affinity purification using a turboGFP antibody (Evrogen) was used to isolate fusion proteins followed by immobilization onto protein G beads. For immunoblots, proteins were resolved by SDS-PAGE using 3-8% Tris-Acetate gels or 4-12% Bis-Tris precast gels (NuPage Life Technologies), then transferred using XCell II Blot module (Life Technologies) to either nitrocellulose or PVDF membranes. Protein bands were detected using SuperSignal West Femto or Pico kit (Thermo Scientific). Table S3 contains a list of reagents used.

### Mass Spectrometry and proteomic data analysis

For whole tissue proteome analysis, the Filter Aided Sample Preparation (FASP) method was used^42^. Briefly, mouse testes samples were homogenized in a lysis buffer consisting of 100mM Tris (hydroxymethyl)amino-methane hydrochloride (Tris-HCl), pH 7.5, in presence of protease (Complete Mini Protease Inhibitor Tablets, Roche and 1mM Phenylmethylsulfonyl fluoride, Sigma) and phosphatase inhibitors (PhosSTOP Phosphatase Inhibitor Cocktail Tablets, Roche). Samples were further processed and peptides and proteins were identified and quantified with the MaxQuant software package, and label-free quantification was performed by MaxLFQ, as described in ^43^. The false discovery rate, determined by searching a reverse database, was set at 0.01 for both peptides and proteins. All bioinformatic analyses were performed with the Perseus software. Intensity values were log-normalized, 0-values were imputed by a normal distribution 1.8 π down of the mean and with a width of 0.2 π. Statistically significant variance between the sample groups was tested by a permutation-based FDR approach and a Student’s t test with a p value cut-off of 0.01. Total proteomic data are available via ProteomeXchange with identifier PXD006849 and are summarized in Table S2.

To examine endogenous ZMYND10 interactions from postnatal day 30 (P30: a period of synchronized flagellogenesis) testes extracts using two well-validated polyclonal antibodies (ZMYND10 Proteintech and Sigma) using an IP/MS workflow carried according to ^44^. Mass spectra were analysed using MaxQuant software and label-free quantification intensity values were obtained for analysis. T-test p-values between MS runs were calculated. MS datasets were ranked by log_2_ fold-change (enrichment) over IgG controls (Table S1). As a filtering strategy to find 'true' interactions, we used the CRAPome repository (http://www.crapome.org/.) containing a comprehensive list of the most abundant contaminants commonly found in AP/MS experiments^45^. To aid filtering, we used an arbitrary threshold of 25 (i.e. proteins appearing in >25 out of 411 experiments captured in the CRAPome repository) were removed from further analysis. Filtered interactors common to both ranked datasets were prioritized for further studies for validation as interactors of ZMYND10 *in vivo*. The mass spectrometry proteomics data have been deposited to the ProteomeXchange Consortium via the PRIDE partner repository with the dataset identifier PXD006849.

### Ependymal primary cultures

Multiciliated ependymal primary cultures were setup by isolating neural stem cells from E18.5 mouse brains according to the protocol^46^. Briefly, a mouse brain was isolated by removing the cranial flaps and further sub-dissecting it in Hank’s medium (HBSS 1 × without Ca^2+^ and Mg^2+^, with 0.075% sodium bicarbonate, 0.01 M HEPES and 1× Pen/Strep medium). The meninges, olfactory bulbs and hindbrain regions were removed. Then, the brain was cut open along the inter-hemispheric fissure and parenchyma surrounding the corpus callosum was removed. A coronal incision was made near the hindbrain to easily remove/peel off the hippocampus along the caudo-rostral axis. The lateral wall of the lateral ventricle that contains the sub-ventricular zone was exposed. Ventricular cup preparations were trypsinized for upto 60 minutes at 37°C using TrypLE express cell dissociation enzyme (Life Technologies). Trypsinized tissue was further manually dissociated and cells were suspended mechanically in DMEM containing 1% Pen/Strep. Cell suspensions were plated onto Laminin coated wells of a 24-multiwell glass bottom plate (Sensoplates from Greiner) and maintained in DMEM-10% FCS (without antibiotic selection) in a humidified 5% CO2 atmosphere at 37°C. Cells were inspected after 5 days to look for morphological changes such as tight junction and hexagonal epithelial monolayer formation. Once cells had formed an epithelial monolayer, cell culture medium was replaced to DMEM with reduced serum (1% FCS) to induce multiciliogenic cell differentiation. Complete cell differentiation into multiciliated cells was achieved after up to 21 days post serum deprivation.

### Reverse transcription Quantitative Real time-PCR (RT qPCR)

Total RNA was isolated from freshly dissected tissue or tissue stored in RNAlater (Qiagen). Isolation was carried out using RNeasy Mini Kit or RNeasy Fibrous Tissue Mini Kit (Qiagen) following manufacturer’s protocol. RNA samples were treated with Turbo DNAse to remove genomic DNA contamination using the Turbo DNA free kit (Ambion). Intron-spanning RT-qPCR assays were designed using the Universal Probe Library probe finder tool (Roche) to identify transcript specific primer-probe sets listed in supplementary table. Three separate experimental runs were carried out for each plate. All runs were done on three individual biological replicates. To calculate relative amounts of transcripts in a sample, standard curves were generated using either serial dilutions of wild-type cDNA. Target Cp were normalised to the Cp values of the reference gene *Tbp* which showed largely invariant expression in wild type and mutant samples. Normalised Cp values were used to extrapolate transcript abundance from the standard curves generated. Data was analysed using Roche LC480 software. Subsequently, a paired two-tailed students t-test was used to compare differences in the mean expression values between wild type and mutant samples.

### DM-CHX FKBP8 inhibitor studies

Lyophilized FKBP8 inhibitor N-(N’N’-Dimethylcarboxamidomethyl)cycloheximide (DM-CHX) ^24^ was dissolved in sterile PBS in a 1mM stock and diluted further to working concentrations in MucilAir™ media (EPITHELIX Sàrl.). MucilAir tracheal epithelial cultures (EPITHELIX Sàrl.) inserts from healthy human donors (same for each stage, immature D19 after air-lift or mature D60 after air-lift) were incubated with DM-CHX for 24 hours and harvested in mild lysis buffer, (50 mM Tris-HCl (pH 7.5), 100 mM NaCl, 10% Glycerol, 0.5 mM EDTA, 0.5% IGEPAL, 0.15% Triton-X 100 and Halt Protease Inhibitor Single use cocktail EDTA free (Thermo Fischer)).

## Acknowledgements

The authors thank IGMM core services, the IGMM imaging and animal facilities for advice and technical assistance. This work was supported by core funding from the MRC (<u>MC_UU_12018/26</u>) (to GRM, PY, MK, PM), and a Science Foundation Ireland grant (SIRG) (to AGM and AvK).

---

**Figure S1.**
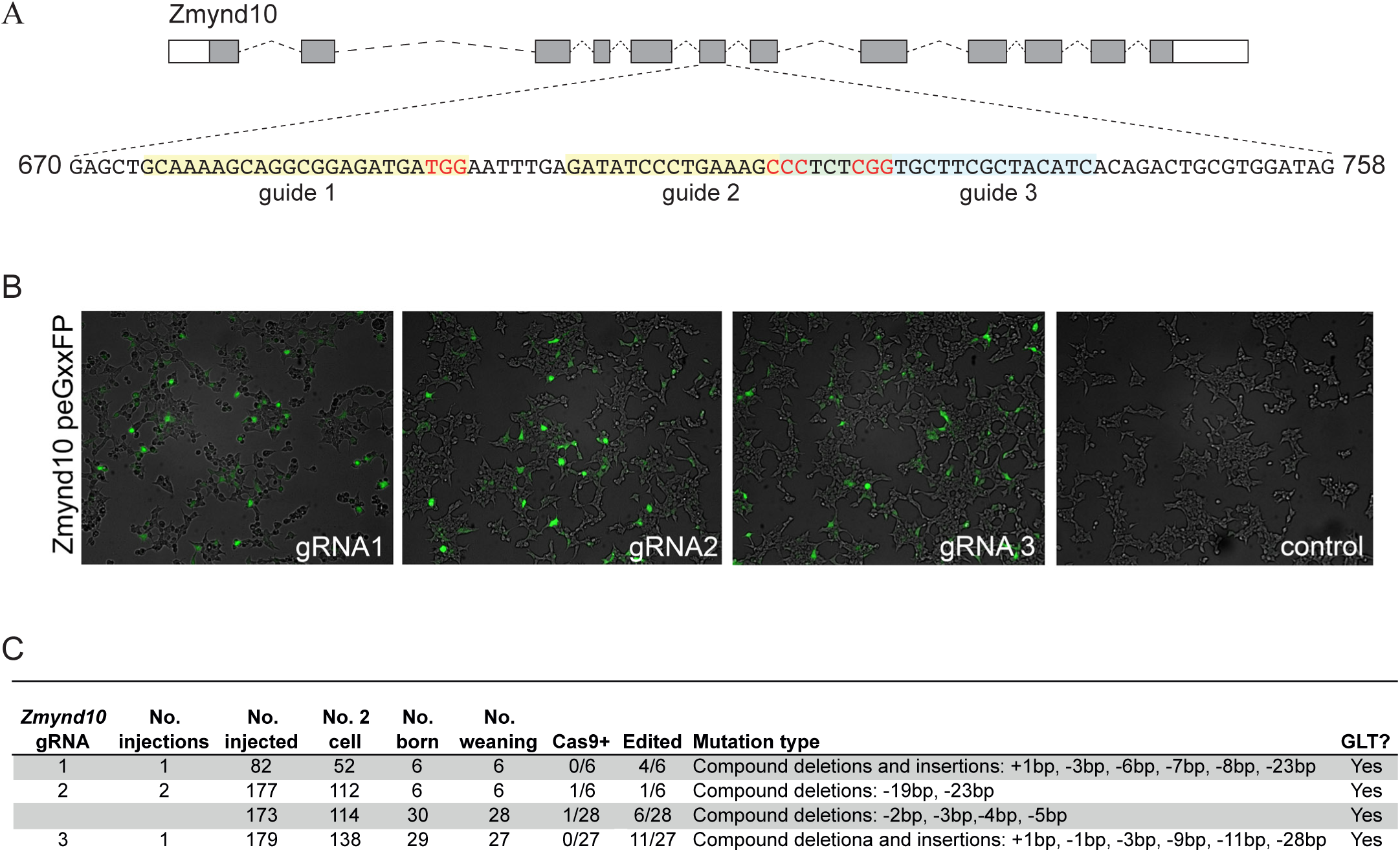
Generation of *Zmynd10* mutant mice by CRISPR gene editing. (A) Schematic of *Zmynd10* mouse locus (ENSEMBL: ENSMUSG00000010044) and guide design targeting critical exon 6 which were cloned into px330 (Addgene:#42230)^38^. (B) Validation of activities of gRNAs to generate double-stranded breaks were initially tested in HEK293 cells by using a reporter to assay reconstitution of EGFP upon cleavage (Addgene: #50716)^39^. (C) Table summarizing pronuclear microinjection rounds for generating *Zmynd10* mutants.

**Figure S2.**
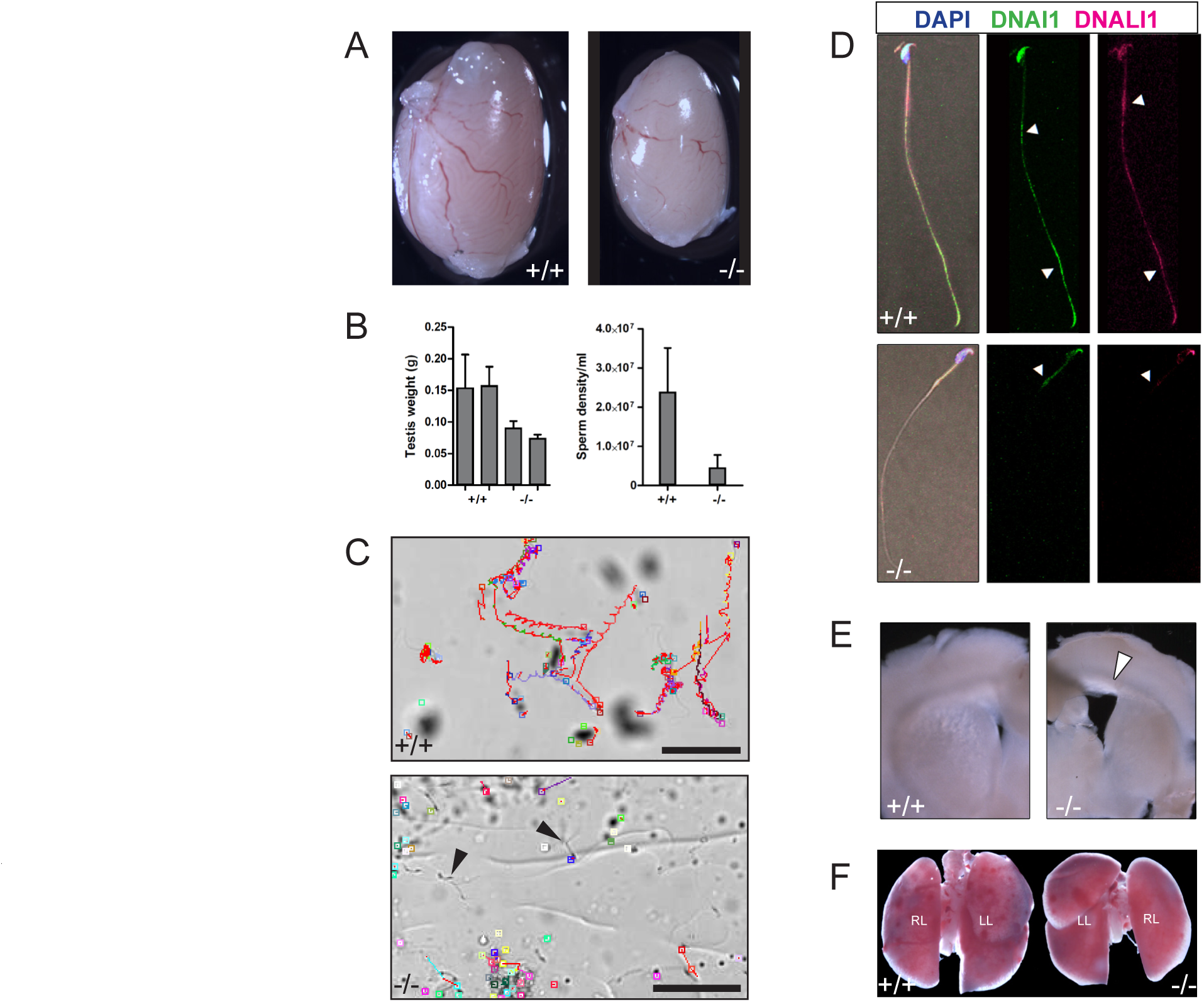
Detailed phenotypic analysis of ☐ZMYND10 mutants. (A) Representative image of a testis from *Zmynd10* mutant and wild type control (P150). (B) As a result of defects in flagellar development, mutants have smaller testes(mean +/-s.e.m, n=3 males/genotype). *Zmynd10* mutants have lower sperm density (mean +/-s.e.m, n=3 males/genotype). (C) *Zmynd10* mutant sperm has reduced motility. Arrows point to static/paralyzed sperm in mutants. Scale bars= 50μm. (D) Immunofluorescent staining of DNAI2 and DNALI1 is absent along *Zmynd*^-/-^ flagella Arrowhead in wild type point to the annulus region and flagellar tip. In the mutant, the arrowhead points to the annulus. (E) Coronal vibratome sections of brains show *Zmynd10* mutants display dilated lateral ventricles (arrowhead) due to hydrocephaly. (F) Representative image of a gross dissection of lungs show *situs inversus totalis* in mutants.

**Figure S3.**
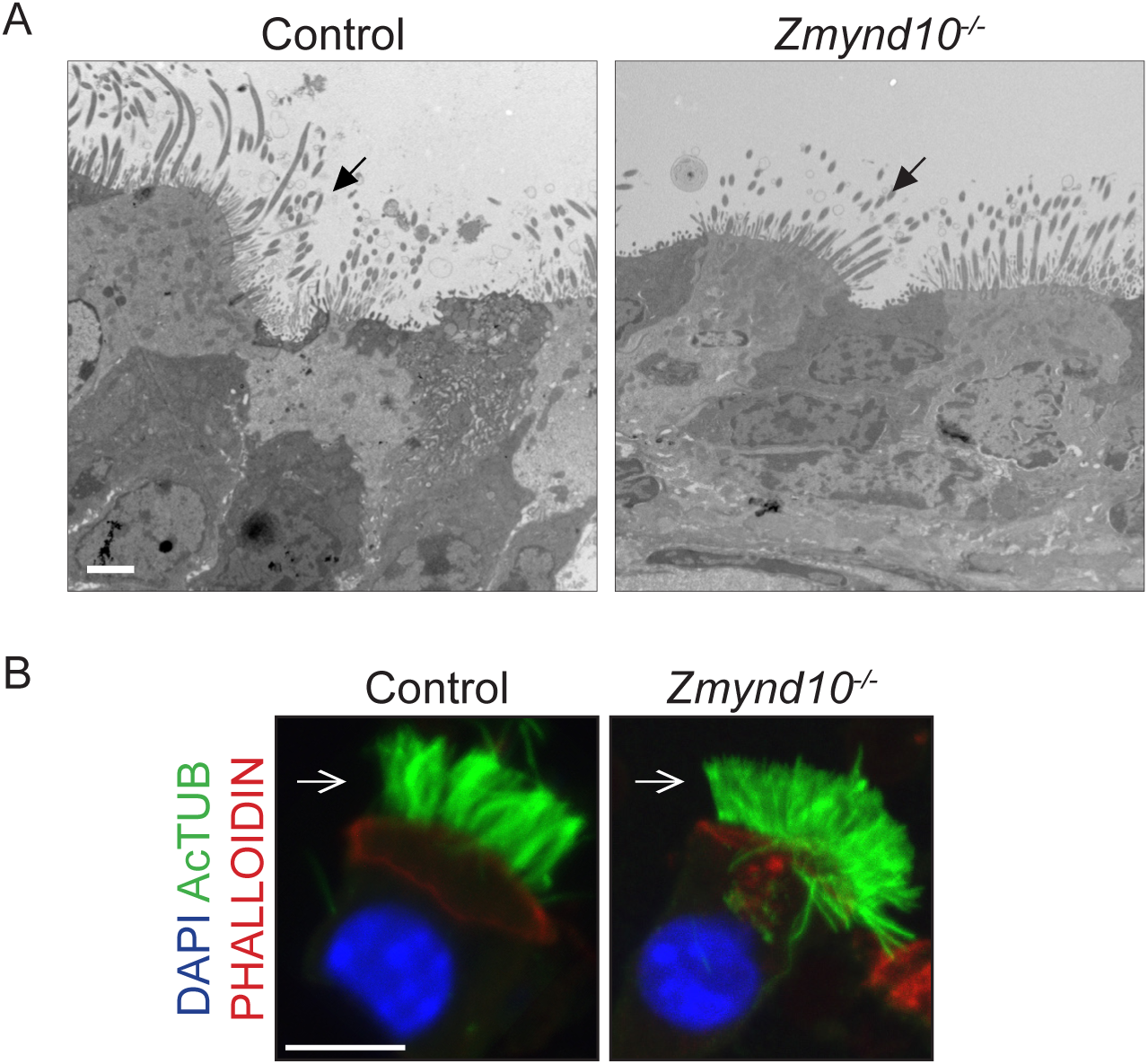
No gross cilial defects in *Zmynd10* mutants. (A) Transmission electron micrographs (TEM) of tracheal multiciliated cells from wild type and mutant female mice (P9). Arrows point to cilia on luminal surface of the cells; basal bodies appear correctly docked in the mutant cells similar to controls. Scale bar = 2μm. (B) Immunostaining for cilia (acetylated α-tubulin) and apical F-actin (phalloidin) reveals no cytoskeletal gross defects in *Zmynd10* mutants. Scale bars = 10μm

**Figure S4.**
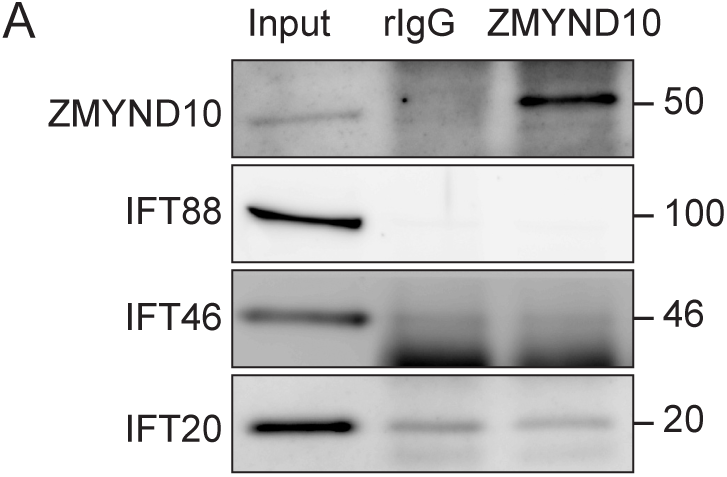
Interactions between ZMYND10 and IFTs or DNAAFs were not detected. (A) ZMYND10 pull-down from fully differentiated multiciliated ependymal cells shows that there is no interaction between ZMYND10 and IFT-B proteins, consistent with a lack of gross ciliary defect.

**Figure S5.**
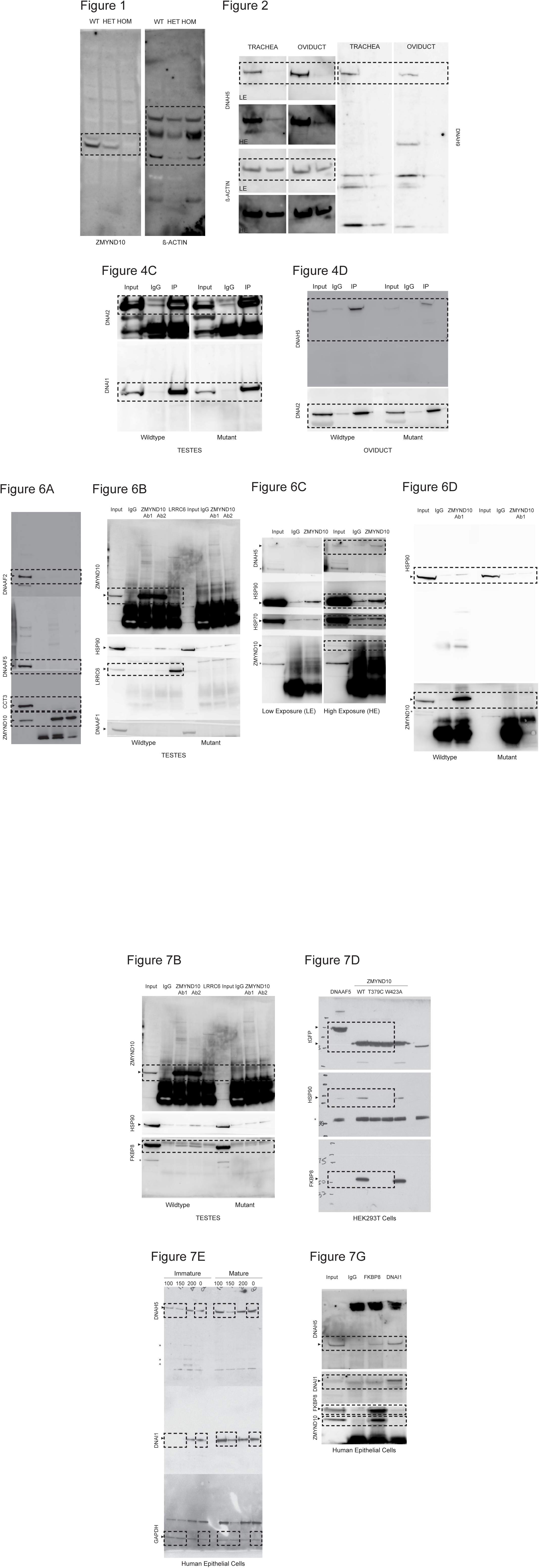
Uncropped immunoblots. Images show uncropped versions of immunoblots. Figure panels used are demarcated with a box and the detected proteins are labeled. Asterisks are used to denote either non-specific bands (6C, 6D and 7B) or putative degradation products (7E). HE= High exposure, LE=Low exposure

**Movie 1: Rapid ependymal ciliary motility in lateral ventricles of a wild-type mouse**

High-speed video microscopy on a coronal brain vibratome section (postnatal day 11 mouse, littermate control) shows ependymal cilia lining lateral ventricles beating with high frequency in a wild-type mouse.

**Movie 2: Immotile ependymal cilia lining lateral ventricles of *Zmynd10* mutant mouse**

High-speed video microscopy on a coronal vibratome section of a brain from a *Zmynd10* mutant mouse (postnatal day 11) shows complete loss of ependymal cilia motility.

**Movie 3: Control murine ependymal cilia with metachronal waveform**

High-speed video microscopy on a coronal vibratome section of a *Zmynd10* mild hypomorphic mutant mouse brain (p. M179del; postnatal day 24) shows arrays of cilia beating in a metachronal waveform and actively generating fluid flow to move particulates over the ventricle tissue.

**Movie 4: Tufts of immotile ependymal cilia in *Zmynd10* mutant murine brain**

High-speed video microscopy on a coronal vibratome section of a *Zmynd10* null mutant mouse brain (p. L188del; postnatal day 29) shows arrays of immotile cilia lining the ventricle tissue with no active fluid flow noticeable.

**Movie 5: Aberrant flagellar motility in ZMYND10 mutant murine epididymal spermatozoa**

High-speed video microscopy on mature spermatozoa extracted from the epididymis of a 5 month old *Zmynd10* CRISPR founder mutant mouse and slowed-down in methylcellulose. The majority of spermatozoa were completely immotile but rarely displayed highly aberrant flagellar movements as observed in the video.

**Movie 6: Sinusoidal flagellar motility in wild type murine epididymal spermatozoa**

High-speed video microscopy on mature epididymal spermatozoa extracted from a 5 monthold wild-type mouse (littermate control) and slowed-down in methylcellulose. Virtually all spermatozoa underwent forward motion with the flagella displaying a sinusoidal beat pattern.

**Supplemental Table 1.**
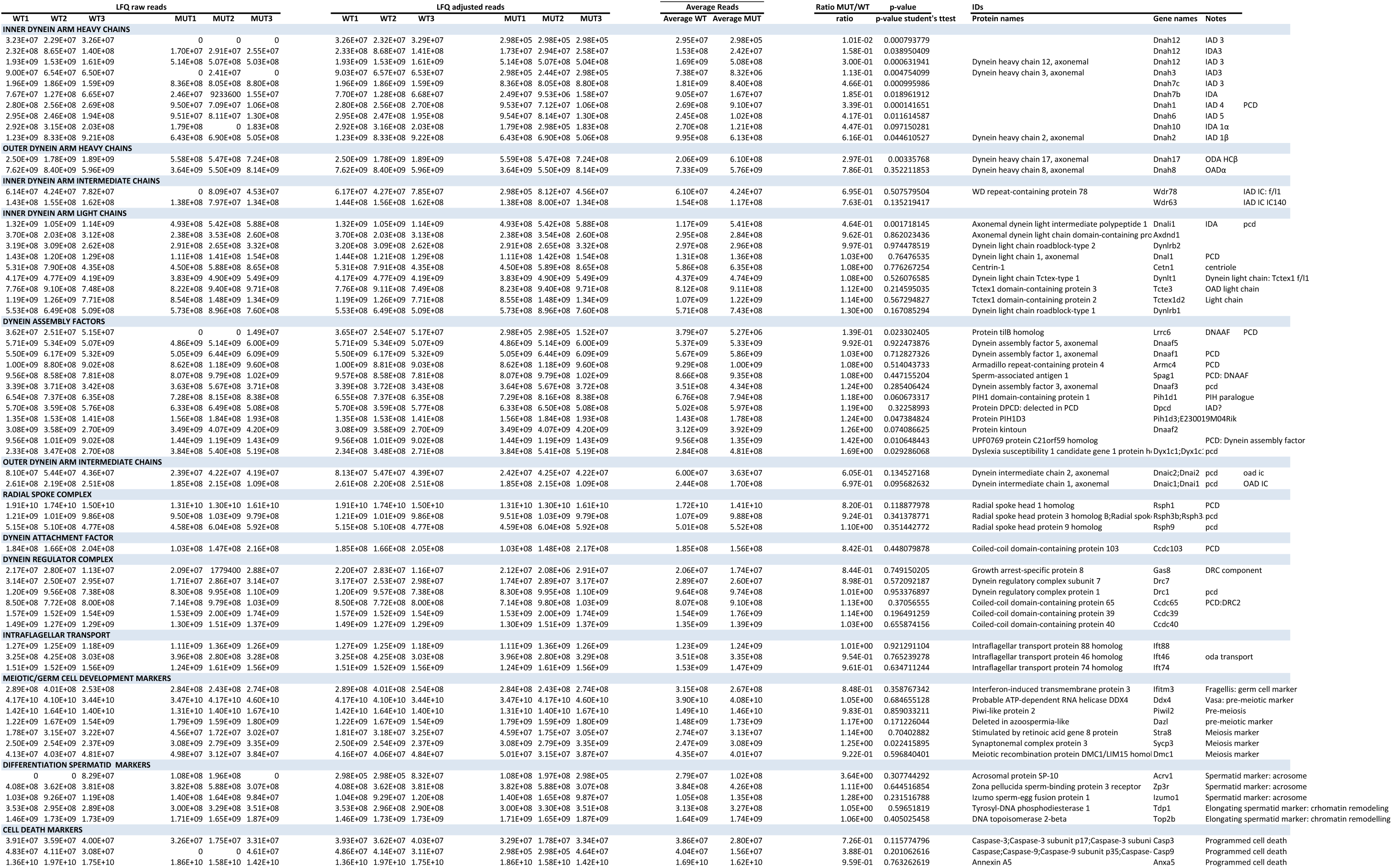
Label-free quantitative global proteomics of P25 testes from Zmynd10+/+ and Zmynd10−/− mice.

**Supplemental Table 2.**
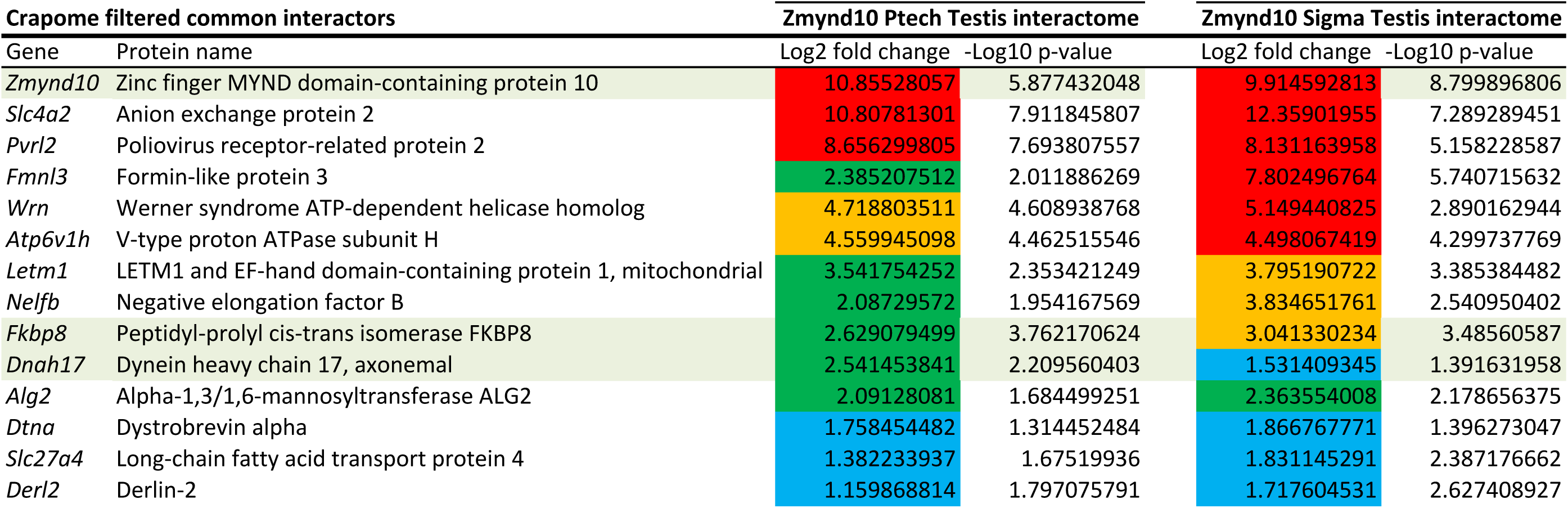
Endogenous ZMYND10 immunoprecipition protein interactor profiling

**Supplemental Table 3.**
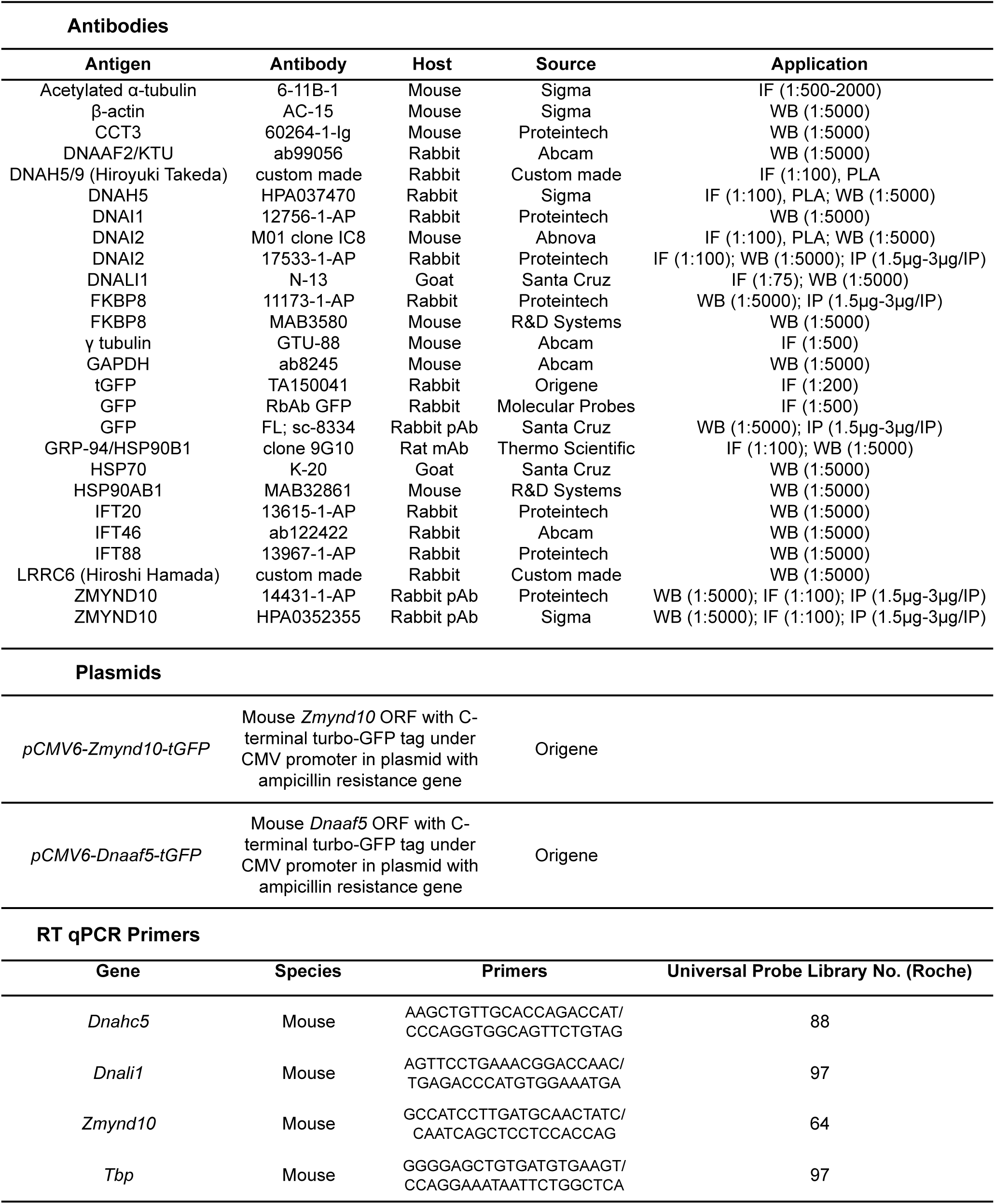
Reagents

